# Atypical and distinct microtubule radial symmetries in the centriole and the axoneme of *Lecudina tuzetae*

**DOI:** 10.1101/2022.02.07.479401

**Authors:** Alexandra Bezler, Alexander Woglar, Fabian Schneider, Friso Douma, Léo Bürgy, Coralie Busso, Pierre Gönczy

## Abstract

The centriole is a minute cylindrical organelle present in a wide range of eukaryotic species. Most centrioles have a signature 9-fold radial symmetry of microtubules that is imparted onto the axoneme of the cilia and flagella they template, with 9 centriolar microtubule doublets growing into 9 axonemal microtubule doublets. There are exceptions to the 9-fold symmetrical arrangement of axonemal microtubules, with lower or higher fold symmetries in some species. In the few cases where this has been examined, such alterations in axonemal symmetries are grounded in likewise alterations in centriolar symmetries. Here, we examine the question of microtubule number continuity between centriole and axoneme in flagellated gametes of the gregarine *Lecudina tuzetae*, which have been reported to exhibit a 6-fold radial symmetry of axonemal microtubules. We used time-lapse differential interference microscopy to identify the stage at which flagellated gametes are present. Thereafter, using electron microscopy and ultrastructure-expansion microscopy coupled to STimulated Emission Depletion (STED) super-resolution imaging, we uncover that a 6- or 5-fold radial symmetry in the axoneme is accompanied by an 8-fold radial symmetry in the centriole. We conclude that there can be plasticity in the transition between centriolar and axonemal microtubules.

## Introduction

Centrioles exhibit a near universal 9-fold radial symmetry of microtubules [reviewed in ^1^]. In ciliated and flagellated cells, centrioles mature into basal bodies, which seed the formation of a likewise 9-fold radially symmetrical axoneme, the inner core of cilia and flagella [reviewed in ^2^]. Thus, the symmetry of the centriole is thought to systematically directly dictate that of the axoneme, but experimental evidence supporting this view is sparse.

Phylogenetic considerations indicate that the near universal 9-fold radial symmetry of centriolar and axonemal microtubules was present in the last eukaryotic common ancestor [reviewed in ^3^]. The strong evolutionary pressure to maintain this architecture is thought to reflect the critical importance of ciliary and flagellar motility across a vast range of organisms, including protists, algae, animals, as well as some plants and fungi. Loss of this conserved arrangement is expected to result in reduced cell motility and thus usually be selected against.

In most organisms, the proximal part of centrioles is characterized by 9 microtubule triplets – dubbed A, B and C microtubules; the A and B microtubules in each triplet extend until the distal end of the organelle. Upon cilium and flagellum formation, these 9 microtubule doublets extend further to form the peripheral microtubule doublets of the axoneme. In most motile cilia and flagella, two centrally located axonemal microtubules are generated in addition using a distinct mechanism [^4^, reviewed in ^5^].

Despite such general stereotypy, structural variations of the centriole have evolved multiple times independently. For instance, microtubule singlets – instead of the usual triplets and doublets-are present in some apicomplexans and fungi, as well as in *Caenorhabditis elegans*, with the singlets being also arranged in a 9-fold radially symmetrical fashion [^6–10^]. Because such centriolar microtubule singlets are followed by axonemal microtubule doublets, in these cases some peripheral axonemal microtubules must be generated by other mechanisms than simply a continuation of centriolar microtubule growth.

There are also instances that deviate from the canonical 9-fold radially symmetrical arrangement of peripheral microtubules in the axoneme; this is usually accompanied by a lack of central microtubules and impaired cell motility. Examples of axonemes with a higher number of microtubule doublets, namely 10, 12, 14, 16 or 18, are encountered in certain protozoans, insects, arthropods and nematodes [^11–16^]. The corresponding centriole symmetry was investigated in two such cases, the Protura *Acerentulus trägardhi* and the sea spider *Nymphon leptocheles* [^14,17^]. In both cases, the axoneme and the centriole exhibit the same 12-fold radial symmetry. An extreme instance occurs in the giant axoneme of the insect *Sciara oprophila*, where up to 70 microtubule doublets can be present [^18,19^]. Also in this case, the radial symmetry of centriolar microtubules mirrors that of the axoneme.

Axonemes can also have radial symmetries of orders lower than 9; for instance, 8-fold symmetrical axonemes are present in the coccidian *Eimeria* sp. [^20^]. A further reduction is encountered in the gregarines *Diplauxis hatti* and *Lecudina tuzetae*, in which 3- and 6-fold radial symmetry of axonemal microtubule doublets have been reported, respectively [^21,22^]. The underlying centriole symmetry has not been investigated in any organism with lower than 9-fold symmetry of axonemal microtubules.

Here, we set out to investigate the above question in the flagellated gametes of *L. tuzetae*, which have been reported to exhibit a 6-fold radial symmetry of axonemal microtubule doublets, with one case of 5 doublets plus 1 singlet also having been observed [^22^]. Using electron microscopy (EM) and ultrastructure-expansion microscopy (U-ExM) coupled to STimulated Emission Depletion (STED) super-resolution imaging of flagellated gametes, we show that axonemal microtubule doublets exhibit a 6- or 5-fold radially symmetrical arrangement. Importantly, in addition, we establish that this is accompanied by an 8-fold radial symmetry of microtubules in the centriole, thus uncovering unprecedented plasticity in the transition between centriolar and axonemal microtubules.

## Results

### *L. tuzetae* flagellated cells harbor the centriolar marker Centrin

Using time-lapse differential interference (DIC) microscopy, we monitored gametogenesis inside the ∼100 μm large *L. tuzetae* gametocyst, which stems from a pair of trophozoite cells that are separated by a membrane and whose progeny nuclei thereby occupy distinct sectors (Fig. 1A; Supplemental Video 1). After several rounds of nuclear divisions inside these two sectors (Fig. 1A, 0 h 00 min), cellularization takes place, such that several hundred of flagellated as well as non-flagellated gametes are present in the gametocyst (Fig. 1A, 5 h 34 min). The onset of flagellar movement is followed by the mixing of the two gamete types, a phase that has been dubbed “the dance of the gametes” [^22,23^], and then fertilization (Fig. 1A, 6 h 13 min). Eventually the gametes stop moving and newly formed zygotes develop within the gametocyst (Fig. 1A, 7 h 31 min).

**Figure 1.**
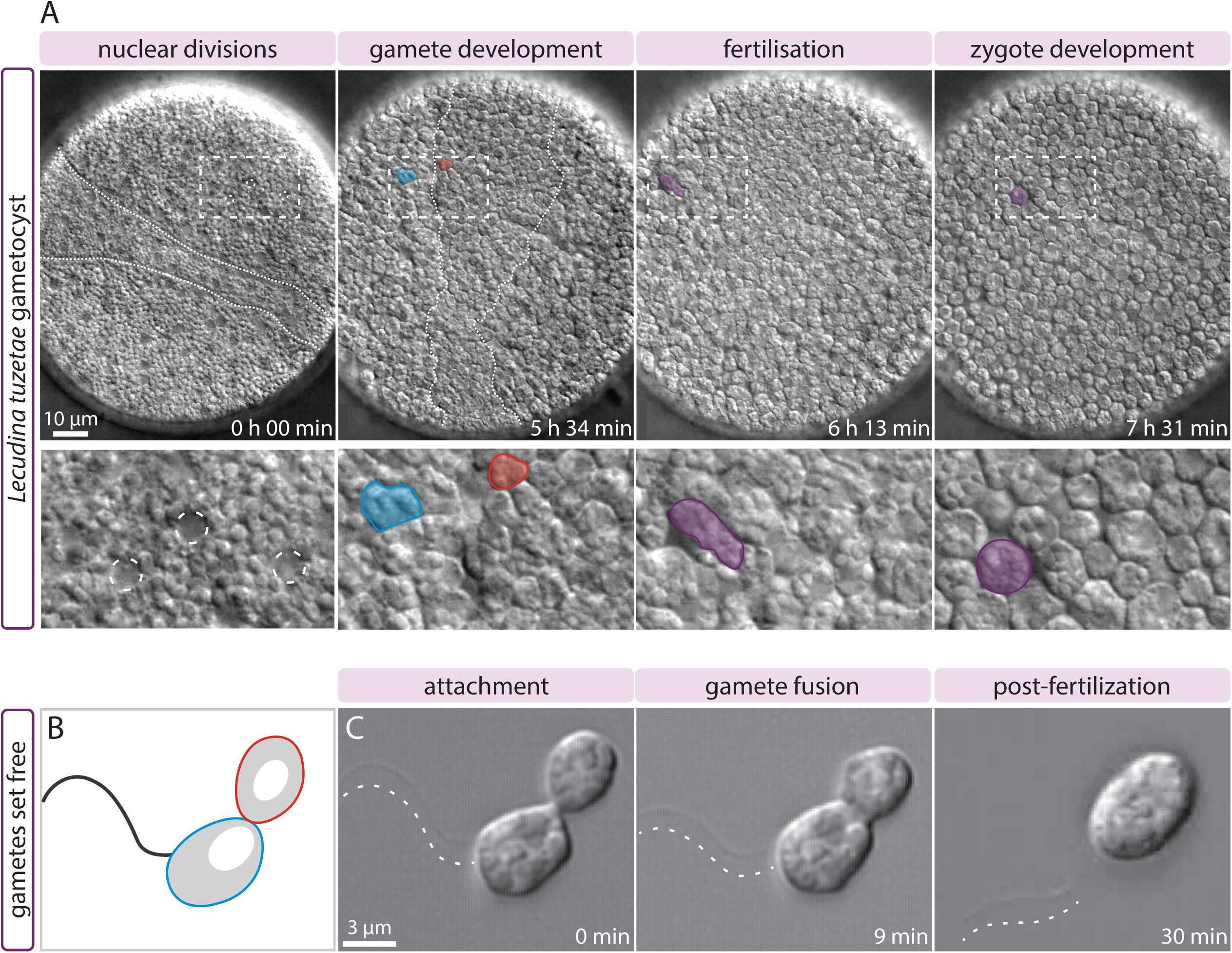
Live imaging of flagellated *L. tuzetae* gametes. (A) Differential interference contrast (DIC) time-lapse microscopy of *L. tuzetae* gametocyst development in distinct sectors (dashed lines), reflecting territories occupied initially by the two parental trophozoites (top: time from start of recording), with two-fold magnified insets (bottom); see also Supplemental Video 1. (0 h 00 min) Nuclei (dashed circles) are visible as areas devoid of granules during the nuclear divisions that occur within the two sectors. (5 h 34 min) Cellularization individualizes nuclei into larger flagellated (blue) and smaller non-flagellated (red) gametes within the two sectors. Note that these two gamete types have also been referred to as male and female gametes, respectively [^22^]. Note also that flagellated and non-flagellated gametes each occupy ∼50% of the gametocyst volume, but that a single focal plane is visible here. (6 h 13 min) Flagellated gametes are thought to generate flows that mix the two gamete types, which eventually fuse during fertilization (purple). (7 h 31 min) After fertilization, the zygote stops swimming and rounds up (purple). (B) Schematic of encounter between a flagellated (blue) and a non-flagellated (red) gamete. (C) DIC time-lapse microscopy of *L. tuzetae* gamete fusion between flagellated and non-flagellated cells (time from start of the recording); see also Supplemental Video 2. (0 min) Attachment: initially a narrow attachment between cells is observed. (9 min) Gamete fusion: the apposed region widens during this process. (30 min) Post-fertilization: the zygote has formed. The dashed line in the three panels highlights the flagellum located just above it.

To distinguish individual flagellated and non-flagellated gametes, as well as to better analyze fertilization events, gametes were set free from the gametocyst (Fig. 1B). Time-lapse DIC microscopy enabled us to observe the movement of the flagellum (Fig. 1C; Supplemental Video 2), as well as to monitor the encounter and the progressive fusion of the two gamete types, which results in zygote formation (Fig. 1C).

As in other gregarines, the axoneme of *L. tuzetae* is comprised of two parts: an internal cytoplasmic segment and an external segment, which is motile [^22^, reviewed in ^24^]. We analyzed flagellated gametes using indirect immunofluorescence microscopy with antibodies against α-tubulin and tyrosinated α-tubulin. Both internal cytoplasmic and external segments could be visualized with these antibodies (Fig. 2A, white outline indicates cell body). We likewise detected the axoneme in flagella extracted with their centrioles (hereafter referred to as extracted flagella) using antibodies against α-tubulin and acetylated tubulin (Fig. 2B). Importantly, antibodies against human Centrin-2, which mark centrioles in a large range of organisms, also localized to the base of extracted flagella, at the center of the microtubule asters (Fig. 2C), where γ-tubulin was shown to localize [^25^]. We conclude that Centrin proteins, and therefore centrioles, are likely present at the base of the axoneme in *L. tuzetae* flagellated gametes.

**Figure 2.**
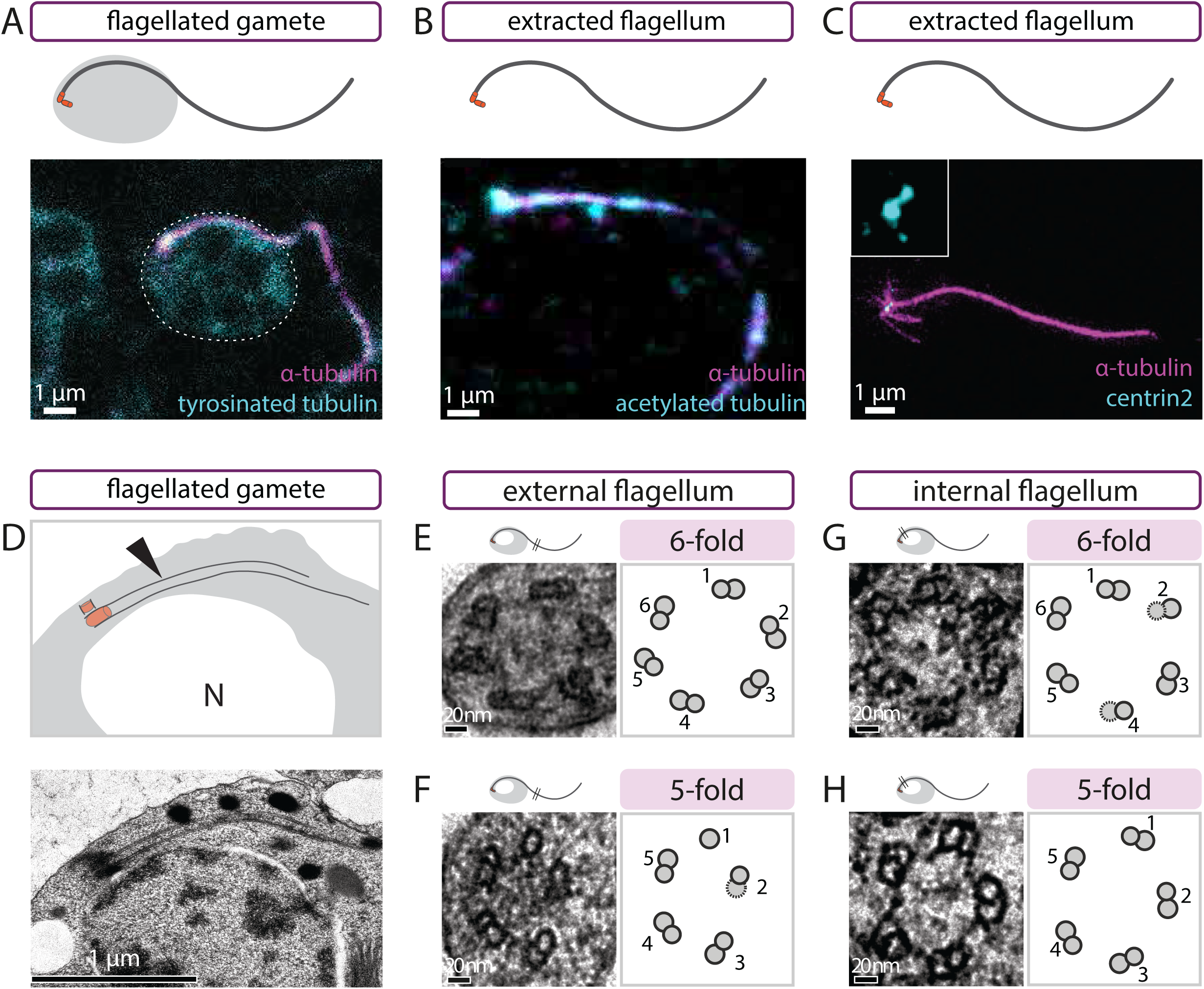
Flagellated gametes harbor Centrin and exhibit 6- or 5-fold symmetry. (A) Schematic (top) and corresponding maximum intensity projection confocal image of *L. tuzetae* flagellated gamete stained with antibodies against α-tubulin (magenta) and tyrosinated-tubulin (cyan). Dashed line indicates the cell body. (B) Schematic (top) and corresponding single plane confocal image of extracted *L. tuzetae* flagellum stained with antibodies against α-tubulin (magenta) and acetylated-tubulin (cyan). (C) Schematic (top) and corresponding maximum intensity projection STED image of extracted *L. tuzetae* flagellum stained with antibodies against α-tubulin (magenta) and Centrin-2 (cyan), with four-fold magnified inset showing two Centrin-2 foci at the base of the flagellum. (D) Schematic (top) and corresponding transmission EM of *L. tuzetae* flagellated gamete with transverse view of centriole and internal axoneme next to the nucleus. Arrowheads points to internal part of the flagellum. (E-H) Transmission EM transversal view of *L. tuzetae* external (E, F) or internal (G, H) axonemal segment with 6-fold (E, G) or 5-fold (F, H) arrangement of microtubule doublets (left) and corresponding schematics (right). Note that the average distance between adjacent A-microtubules is similar in both types of arrangements (6-fold: 54 nm (+/- 8 nm SD), 5-fold: 51 nm (+/-7 nm SD)).

### Axonemal microtubules exhibit 6- or 5-fold radial symmetry

To confirm the ultrastructure of the axoneme in *L. tuzetae* flagellated gametes, we conducted EM of resin-embedded specimens. As for the immunostainings, the internal flagellum was observed by EM to emanate from the basal body and to extend along the nuclear envelope (Fig. 2D). To analyze the fold radial symmetry of microtubules, we focused on transverse views of axonemes. This enabled us to detect in some cases a 6-fold radial symmetrical arrangement of microtubule doublets in the external segment of the axoneme, without any central microtubule, in line with previous findings (Fig. 2E, N=3) [^22^]. Moreover, we observed a 5-fold symmetrical arrangement of microtubule doublets in the external axonemal segment in other cases (Fig. 2F, N=13). In addition, when examining the internal segment of the axoneme, we likewise found both 6- and 5-fold symmetries (Fig. 2G, 6-fold N=2; Fig. 2H 5-fold N=4). We sought to confirm such 6- and 5-fold symmetrical arrangements through radial image symmetrization analysis, which is a classical approach to determine the fold-symmetry of objects such as centrioles and axonemes [^15,26,27^]. As shown in Supplemental Figure S1A-S1B, by applying 4- to 11-fold symmetrization, we found that 6- and 5-, respectively, fold symmetrization bears the most resemblance with the arrangement of microtubules in the original images. We noted also that flagellated gametes with either 6-fold or 5-fold symmetry were present in the same gametocyst, demonstrating that difference between the two types does not reflect genetic heterogeneity between gametocysts. Overall, we conclude that all internal and external segments of *L. tuzetae* axonemes analyzed deviate from the canonical 9-fold symmetry.

### Widening of the microtubule-bearing area towards the centriole

Further inspection of rare longitudinal views of resin-embedded flagellated gametes analyzed by EM uncovered a widening of the area corresponding to microtubules in the region where the centriole is located (Fig. 3A, compare blue and red lines, N=3). Tomographic reconstruction of such longitudinal EM sections confirmed a widening of the microtubule wall in the centriolar region (Fig. 3B, compare blue and red lines, Fig. 3C, N=2). Interestingly, in addition, this analysis indicated that whereas some of the microtubules from the centriole appear to be continuous with those of the axoneme, others are not (Fig. 3C, arrowhead; Supplemental Video 3).

**Figure 3.**
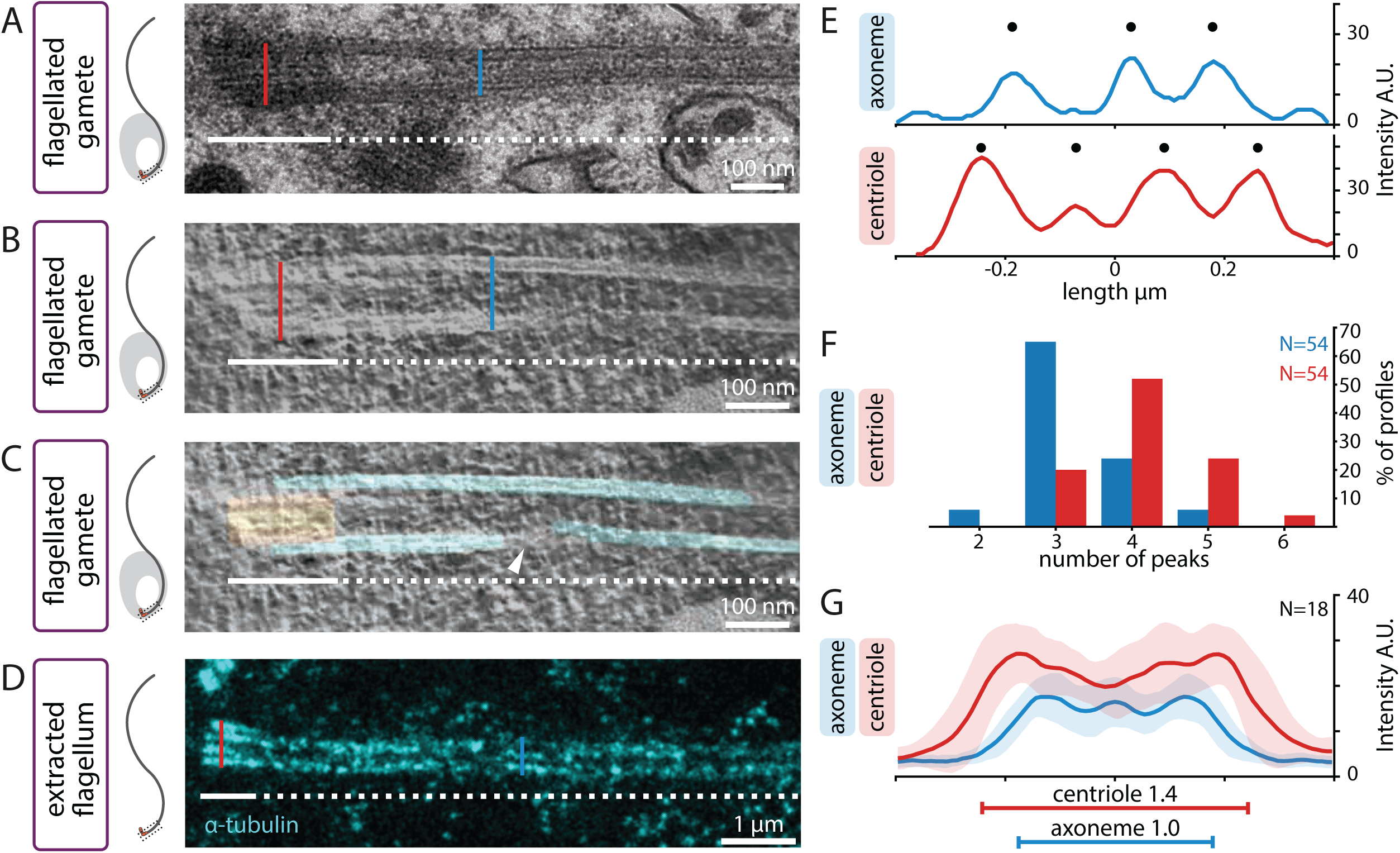
Centrioles are wider than axonemes in *L. tuzetae*. (A) Schematic (left) and corresponding side view transmission EM of *L. tuzetae* internal flagellum embedded in resin after chemical fixation, with centriolar part (white line) and axonemal part (white dashed line). Width of centriole (red line) and axoneme (blue line) are indicated for illustration purposes here, as well as in B and D. (B, C) Side view electron tomogram of 50 nm section embedded in resin after cryo-fixation (B) and corresponding model superimposition (C, microtubules in green, centriolar region in cyan) of *L. tuzetae* internal flagellum, with centriolar part (white line) and axonemal part (white dashed line). Arrowhead points to discontinuity in one of the microtubules. (D) Side view U-ExM STED image from extracted *L. tuzetae* flagellum containing centriole (white line) and axoneme (white dashed line) upon 5.2-fold expansion and after staining with antibodies against α-tubulin. (E) Representative intensity plot profiles measured at centriole (red) and axoneme (blue) from extracted *L. tuzetae* flagellum upon U-ExM STED (from D). In the axoneme, the full-width-half-maximum (FWHM) is 0.44 µm and three intensity peaks (discs) are visible. In the corresponding centriole, the FWHM is 0.58 µm and four intensity peaks (discs) are visible. (F) Line profile peak analysis from population of intensity plot profiles measured at axoneme (blue) and centriole (red). (G) Mean intensity plot profiles measured in three positions for each centriole (red) and axoneme (blue) (N=18 images), with corresponding standard deviations (shaded) from extracted *L. tuzetae* flagella upon U-ExM STED, (N=54 line scans per category). The mean widths at FWHM of axoneme and centriole are statistically different (two-sample Student’s T-test p= 1.22 × 10^−7^) and are indicated by horizontal lines (axoneme: blue, centriole: red).

To quantify the apparent width difference uncovered through EM, we set out to analyze extracted flagella using U-ExM [^28,29^]. We reasoned that the ∼5.2 expansion factor afforded by U-ExM, coupled with STED super-resolution, should enable us to readily probe potential width differences in longitudinal views by immunostaining. U-ExM of such specimens stained with α-tubulin antibodies indeed confirmed the widening towards the centriole (Fig. 3D, compare blue and red lines). Line scans perpendicular to the flagellum revealed that 3 peaks of α-tubulin signal were present in most cases for the axonemal part, irrespective of the peak calling threshold applied, presumably corresponding to the side view of 6- and 5-fold microtubule doublets (Fig. 3E, Fig. 3F blue). By contrast, 4 intensity profile peaks were most frequently observed for the centriolar part (Fig. 3E, Fig. 3F, red). Importantly, measuring the full width half maximum (FWHM) of the centriolar and axonemal line profiles revealed that the width at the centriole is on average ∼1.4 that of the width at the axoneme (Fig. 3G; N=18). Overall, we conclude that the width of the region corresponding to microtubules is consistently larger at the level of the centriole compared to the axoneme emanating from it.

### Centriolar microtubules exhibit 8-fold radial symmetry

The above observations are compatible with at least two possibilities. First, microtubules with 6- or 5-fold radial symmetry could also be present in the centriole but be positioned at a greater distance from one another than in the axoneme, resulting in a larger overall width. Second, the centriole may contain a higher number of peripheral microtubules than the 6 or 5 sets present in the axoneme. Analyzing transverse sections of centrioles should enable one to distinguish between these possibilities. Therefore, we conducted U-ExM followed by STED of extracted flagella probed with α-tubulin antibodies. The rare transverse centriolar views obtained clearly showed more peripheral microtubules than the 6 or 5 that are present in axonemes (see below, N=4). We set out to uncover the symmetry of these centriolar microtubules. To compensate for slight distortions, images of deconvolved transverse views were first circularized (Fig. 4A), before applying symmetrization from 4- to 11-fold, like it was done for the axoneme. As is apparent from Fig. 4B, 8-fold radial symmetrization yielded the best resemblance with the arrangement of microtubules in the original images of the centriole (see also Supplemental Figure S1D and Supplemental Figure S2A). This conclusion was further supported by the 4-fold symmetrization pattern, as expected from an 8-fold symmetrical arrangement (Fig. 4B). For all four centrioles analyzed in this manner, 8- and 4-fold symmetrization correlated best with the original images (Supplemental Figure S2B-C). All other applied symmetries applied resulted in unresolvable microtubule signals that did not resemble the original images (Fig. 4B).

**Figure 4.**
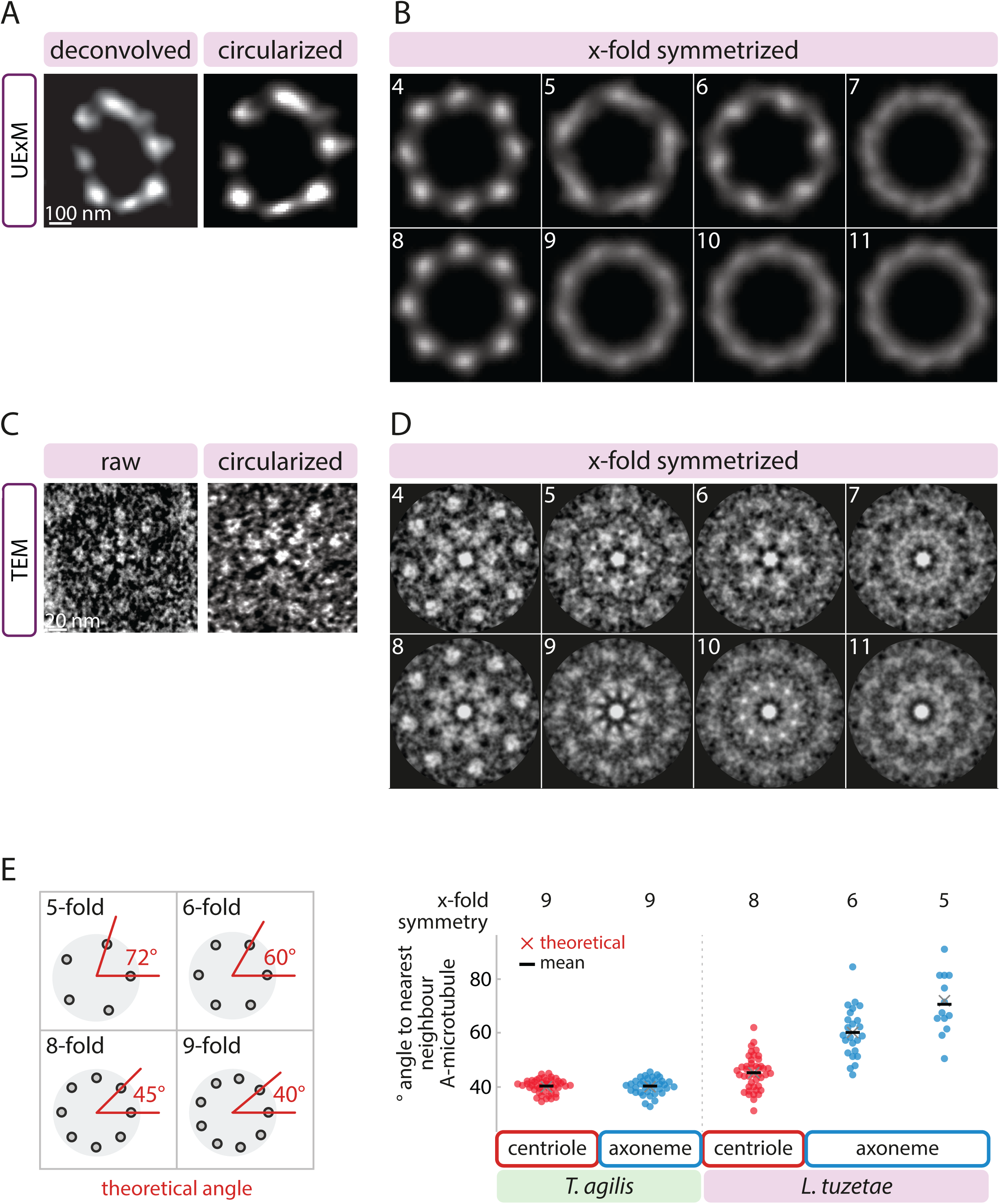
*L. tuzetae* centrioles exhibit 8-fold radial symmetry. (A) Transverse view of deconvolved U-ExM STED image of *L. tuzetae* centriole from extracted flagellum stained with antibodies against α-tubulin (left), and corresponding circularized image (right). (B) X-fold symmetrization (4-11, as indicated) of circularized centriole from (A). (C) Transverse view of transmission EM of *L. tuzetae* centriole embedded in resin after chemical fixation (left), and corresponding circularized image (right). (D) X-fold symmetrization (4-11, as indicated) of circularized centriole from (C). (E) Schematic of theoretical angles in wedges between centriole center and two neighboring peripheral A-microtubules for the indicated fold-symmetries (left). Theoretical angles (red cross) and corresponding measurement of observed angles (disks) with mean (black bar) in *Trichonympha agilis* 9-fold symmetrical centriole (N=45 angles between neighboring microtubules of n=5 centrioles) and axoneme (N=36, n=4 axonemes), as well as in *L. tuzetae* centriole (N=43, n=9) and axoneme (6-fold: N=24, n=4; 5-fold: N=13, n=3), as indicated.

To probe in a complementary manner the radial symmetry of centriolar microtubules, we also analyzed transverse EM sections of the *L. tuzetae* centriole. Not all microtubules could be clearly discerned in a single section in these cases (Fig. 4C), likely because the sections are not perfectly orthogonal to the longitudinal axis of the centriole. Regardless, those centriolar microtubules that were clearly discernable appeared as singlets arranged around a central hub, although we cannot exclude that the electron dense cloud surrounding centrioles or microtubule inner proteins might have obscured B-microtubules. Applying again 4- to 11-fold symmetrization of the EM images, we uncovered that the radial distribution of centriolar microtubules was most compatible with an 8-fold symmetry, a view further supported by the 4-fold symmetrization data (Fig. 4D). Again, for all 8 images analyzed in this manner, 8- and 4-fold symmetrization correlated best with the original images (Supplemental Figure S1C, Supplemental Figure S2D-F). To corroborate this point further in the non-symmetrized raw data, we measured the inner angle in wedges formed between the center of the centriole and two clearly detectable neighboring peripheral microtubules (Fig. 4E). Considering the 360° present in a circle, a given fold symmetrical arrangement of microtubules yields a predictable angle in such wedges. Thus, an 8-fold symmetrical arrangement of microtubules results in a theoretical angle of 45^°^, whereas a 9-fold arrangement yields a theoretical angle of 40^°^. For instance, in an extant TEM data set from *Trichonympha agilis* [^30^], in which microtubules exhibit a canonical 9-fold radially symmetrical arrangement in both centriole and axoneme, the average angle is ∼ 40° in both compartments (centriole: 40.1° +/- 2.5° standard deviation (SD); axoneme: 40.0° +/- 3.0° SD; Fig. 4E). In *L. tuzetae*, by contrast, the average angle measured for all centrioles taken together is 45.1° (+/- 6.3° SD; Fig. 4E), which is fully compatible with an 8-fold symmetrical arrangement. In addition, the average angle between neighboring microtubules within individual centrioles was invariably closer to 45^°^ than to 40^°^ (N=9 centrioles). As anticipated, cross-sections of the 6- and 5-fold axonemes yielded average angles of 60.1° (+/-9.4° SD) and ∼70.6° (+/-11.2° SD), respectively, close to the theoretical angles (Fig. 4E). Together, these findings lead us to conclude that the centriole in *L. tuzetae* exhibits an 8-fold symmetrical arrangement of microtubules underlying 6- and 5-fold axonemes.

## Discussion

Our findings further establish that the *Lecudina tuzetae* axoneme diverges from the canonical 9-fold radial arrangement observed in most species and uncovers the architecture of the corresponding centriole (Fig. 5). We find that the axoneme of the flagellated gamete harbors 6 or 5 radially arranged sets of microtubules, in both internal and external segments of the axoneme, instead of the usual 9. Importantly, in addition, our findings indicate that the centriole from which this axoneme emanates contains 8 microtubules, instead of the usual 9 present in most other species, or of the 6 or 5 that might have been anticipated from the axonemal configuration in *L. tuzetae*.

**Figure 5.**
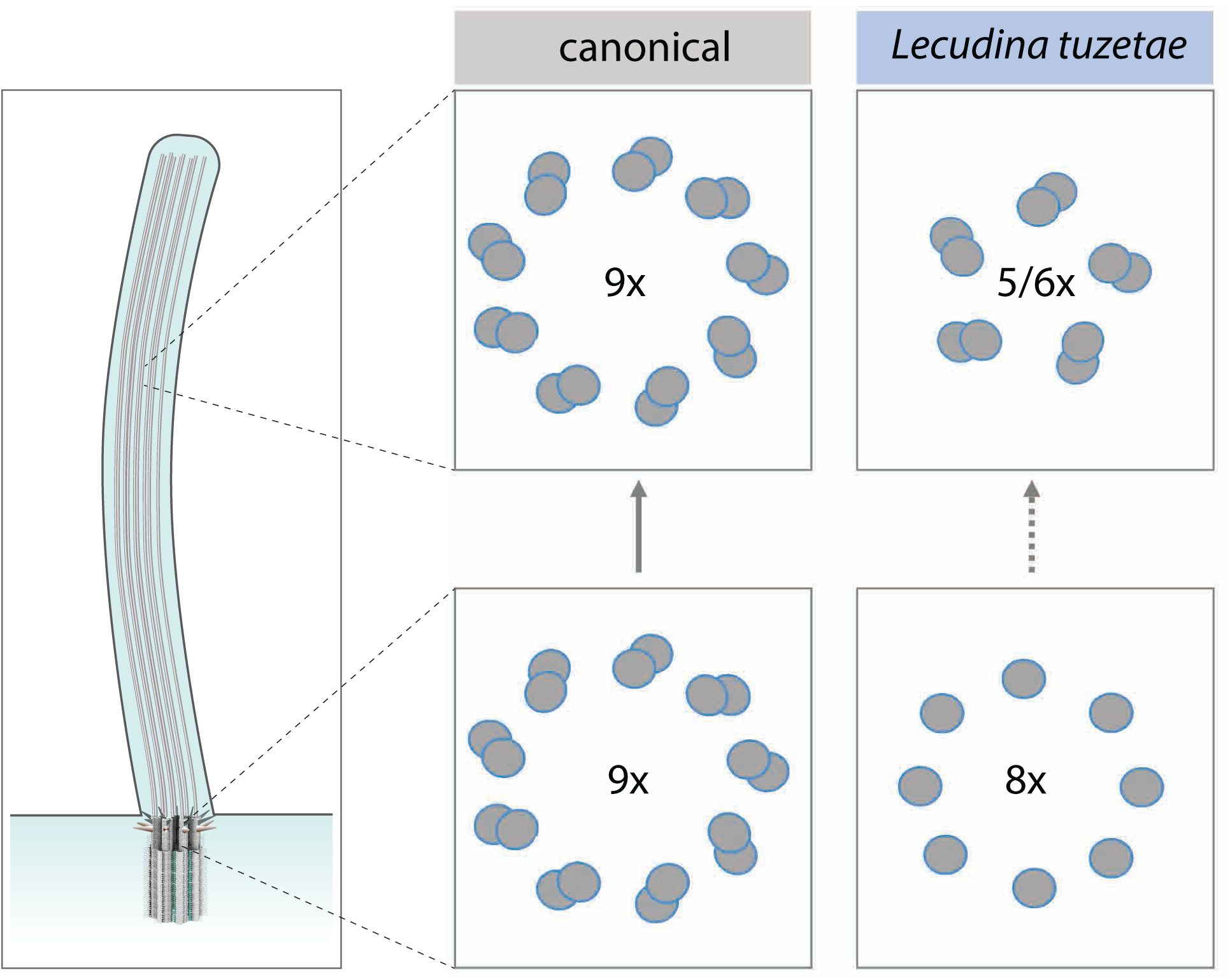
Schematic summary of centriolar and axonemal microtubules in species with canonical 9-fold symmetry and in *L. tuzetae*. Schematic of a flagellum (left). Schematic of microtubule arrangement in the canonical situation, where 9-fold symmetrical microtubule doublets in the centriole yields a likewise 9-fold symmetrical arrangement in the axoneme, as well as in *L. tuzetae*, where 8-fold symmetrical microtubule singlets in the centriole are somehow followed by 5- or 6-fold symmetrical microtubule doublets in the axoneme.

### How can an 8-fold radially symmetrical centriole be built?

What mechanisms could explain the formation of 8-fold radially symmetrical centrioles in *L. tuzetae*? Across the eukaryotic tree of life, proteins of the SAS-6 family are critical for scaffolding the onset of centriole assembly [reviewed in ^31,32^]. SAS-6 forms homodimers, which possess an intrinsic ability to assemble into 9-fold symmetrical ring structures. These ring structures then stack to form a ∼100 nm high cartwheel, which is thought to act as a scaffold for centriole assembly, including by contributing to impart the fold symmetry of peripheral microtubules.

How could *L. tuzetae* SAS-6 (LtSAS-6) self-assembly yield an 8-fold symmetrical structure? In the absence of molecular information regarding LtSAS-6, we speculate that the protein might harbor amino acid changes that favor assembly of an 8-fold symmetrical structure. Compatible with this notion, rational mutagenesis of *Chlamydomonas reinhardtii* SAS-6 (CrSAS-6) can alter the fold symmetry of structures self-assembled *in vitro*, including towards 8-fold [^33^]. Another possibility is based on the finding that although wild-type CrSAS-6 favors self-assembly of 9-fold symmetrical structures, *in vitro*, 8-fold and 10-fold symmetrical ring-structures can be observed in addition to 9-fold symmetrical ones [^33,34^]. Such observations have led to the suggestion that peripheral elements must function together with SAS-6 ring-containing structures to select strictly 9-fold radially symmetrical entities [^33,35^]. Perhaps such peripheral components are absent or different than in most other species in *L. tuzetae*, thus favoring 8-fold symmetrical assemblies.

### Diminution of microtubule numbers from centriole to axoneme

The finding that there is a distinct fold symmetry in the centriole and in the axoneme that emanates from it is unprecedented to our knowledge. It appears that 2 or 3 of the microtubules present in the centriole do not extend further into axonemal microtubules in *L. tuzetae*, indicative of plasticity in the processes at the transition between the two cellular compartments. Interestingly, axonemes assemble rapidly in the cytoplasm of the related apicomplexan *Plasmodium falciparum* [reviewed in ^8^]; perhaps likewise rapid assembly kinetics occur in *L. tuzetae* and may contribute to a plastic transition between centriole and axonemal compartments. Axoneme architecture can also exhibit plasticity over time, as evidenced by specific sensory cilia in *C. elegans*, where the number of microtubule doublets is remodeled from 9-fold in larvae to 6-fold in adults [^36^]. Such temporal remodeling could conceivably also explain why some axonemes exhibit 6-fold and others 5-fold symmetry in *L. tuzetae*.

How could an otherwise very constrained architecture be relaxed in *L. tuzetae* and other apicomplexans? A plausible answer lies in the underlying biology. In contrast to the situation in most other species, the flagellated cells in *L. tuzetae* are enclosed with the non-flagellated cells in a common gametocyst, thus facilitating fertilization. Moreover, since all flagellated cells derive from a single trophozoite, they are likely in competition only with their genetically identical clones. Therefore, the evolutionary pressure for flagellar motility may be minimal in *L. tuzetae*. Compatible with this view, *L. tuzetae* flagellated gametes exhibit a slow tumbling forward motion and are poor swimmers compared to sperm cells from other species (^22,37^). Regardless, such diversity in axonemal and centriolar architecture highlights striking biodiversity in otherwise strictly constrained cellular structures of critical importance for the successful reproduction of many species.

## Acknowledgements

We are grateful to Paul Guichard for contributing to initiate the project, as well as to Isabelle Florent, Graham Knott, Joseph Schrével and Gérard Prensier for precious advice and discussions. Michel Bornens and Gaia Pigino are acknowledged for further discussions. We thank Marie Croisier, Anaëlle Dubois from the Bio-EM Facility in the School of Life Sciences at EPFL headed by Graham Knott for help with ultrastructural analysis, as well as Sophie Booker and Ronan Garnier from the Station Biologique de Roscoff (France) for collecting *Hediste diversicolor* animals. We are grateful to Georgios Hatzopoulos for help with drawing the schematics in Fig. 5. We acknowledge Niccolò Banterle and Ella L. Müller for help with data processing, as well as Isabelle Florent, Paul Guichard, Nils Kalbfuss and Joseph Schrével for useful comments on the manuscript. This work was supported in part by grants from the European Union (MCSA-IF 588594 to AW and COFUND-EuroPostdoc 588459 to FS, as well as ERC AdG 340227 and 588437 to PG).

## Author contributions

AB, AW and PG designed the project. Data was acquired primarily by AB and AW, as well as by FS and CB. AB, AW, FS, FD and LB analyzed data. PG provided supervision. AB, AW and PG wrote the manuscript, with input from all authors.

## Supplemental data

**Video 1 *L. tuzetae* gametocyst development**

Differential interference contrast (DIC) time-lapse microscopy of *L. tuzetae* gametocyst development (time from start of recording). Images were acquired every 10 seconds and are played with 300 times acceleration. Related to Figure 1A.

**Video 2 *L. tuzetae* gamete fusion**

DIC time-lapse microscopy of gamete fusion (time from start of recording). *L. tuzetae* gametes were set free from the gametocyst when they started moving vigorously; arrowhead marks actively beating flagellum. Cells were immobilized on the glass and monitored until zygote formation. Images were acquired every second, registered in FIJI and are played with 10 times acceleration. Related to Figure 1C.

**Video 3 Tomogram with model of microtubules**

Movie rendered in Chimera of the tomographic data from *L. tuzetae* with superimposed model generated in IMOD. The video shows individual slices of the reconstructed tomogram of a 50-nm thick section through a *L. tuzetae* flagellum parallel to the section. Individual microtubules are represented with cylindric models in cyan and the centriole in orange.

**Supplemental Figure S1 Symmetrization of axoneme and centriole to reveal the underlying symmetry**

(A-B) Transmission EM transversal view of *L. tuzetae* axoneme with 6-fold (A; see Figure 2G) or 5-fold (B; see Figure 2H) arrangement of microtubule doublets (left) and X-fold symmetrization (4-11, as indicated; right). Green squares indicate the fold-symmetrization with the highest correlation to the original image, which is 6- and 5-fold, respectively.

(C) Transmission EM transversal view of *L. tuzetae* centriole with unclear arrangement of microtubules (left) and X-fold symmetrization (4-11, as indicated; right). Green squares indicate the fold symmetrization with the highest correlation to the original image, which is 8-fold.

(D) Transverse view of deconvolved U-ExM STED image of *L. tuzetae* centriole from extracted flagellum stained with antibodies against α-tubulin (left) and X-fold symmetrization (4-11, as indicated; right). Green squares indicate the fold symmetrization with the highest correlation to the original image, which is 8-fold.

**Supplemental Figure S2 Symmetrization of the centriole reveals 8-fold symmetry**

(A) Deconvolved STED image of *L. tuzetae* centriole of an extracted flagellum, expanded ∼5.2 x and stained for α-tubulin. Distortions due to the orientation of the centriole were corrected using the ImageJ plugin “Transform-Interactive Affine” (top). Overlays of circularized input image (green) and corresponding 4- to 11-fold symmetrization (magenta) (bottom). Numbers in the top left corner indicate fold symmetry applied in each case.

(B) Corresponding overlays of intensity profiles measured in a circular band along the centriole from circularized (green) and symmetrized (magenta) images.

(C) Box plots of R^2^ values from correlation of intensity profiles between circularized and symmetrized images (N= 4 centrioles). Note that in each case, 8- and 4-fold symmetrized images exhibit the strongest correlation with the input data.

(D) Overlays of circularized input TEM image of a centriole in transverse view (green) and corresponding 4- to 11-fold symmetrization (magenta).

(E) Corresponding overlays of intensity profiles measured in a circular band along the centriole from circularized (green) and symmetrized (magenta) images.

(F) Box plots of R^2^ values from correlation of intensity profiles from input with symmetrized images (N=8 centrioles). Note that in each case, 8- and 4-fold symmetrized images exhibit the strongest correlation with the input data.

## Material and Methods

Work in the laboratory of PG is conducted under authorization number A1821681 2 delivered by the Swiss Federal Office for Public Health.

### Live imaging of *Lecudina tuzetae* gametocysts

*Hediste diversicolor* worms, which host *Lecudina tuzetae* intestinal parasites, were collected in 2017, 2018 and 2021 between April and October from 48°37’8.20’’N 3°57’9.03’W by the collection service of the Station Biologique de Roscoff (France). After courier shipping, worms were maintained without feeding in sea water (22 g/L sea salt) at room temperature, one per 10 cm Petri dish. The Petri dish was aerated on the first day and worms were allowed to recover for 48 h after shipping before being transferred into fresh sea water. This enabled collection of partially synchronous *L. tuzetae* gametocysts during the following 16 h - 20 h from the Petri dish with a pipette (200 µl tip) under a dissecting microscope with transmitted light.

For time-lapse imaging, gametocysts were mounted on 2% agarose pads immersed in sea water, covered with a coverslip, the edges of which were sealed with VaLaP (1:1:1 mixture of petroleum jelly:lanolin:paraffin wax) to prevent evaporation (see Fig. 1A). To mechanically set free flagellated and non-flagellated gametes, gametocysts were mounted without agarose and pressure applied on the coverslip during the “dance of gametes” stage (Fig. 1C). DIC time-lapse microscopy was performed by capturing an image every 10 s for gametocysts and every 1 s for released gametes with a 63x EC Plan-NEOFLUAR objective (NA 1.25) on a Zeiss Axioskop2 plus equipped with a DCC1545M-GL Thorlabs camera. Gametocysts were monitored using DIC time-lapse microscopy to identify the appropriate stage for fixation for subsequent EM and immunofluorescence experiments.

### Immunostaining

For immunostaining of intact gametes and of extracted flagella, gametes were mechanically set free from gametocysts by applying gentle pressure on the coverslip and removing excess water with a filter paper, followed by freeze-cracking on a poly-D-Lysine (2 mg/ml in water) coated slide. To isolate flagella, cysts were collected by centrifugation for 1 min at 100 g on a tabletop centrifuge and lysed in 10 mM K-Pipes pH 6.8, 1% NP-40 for 10 min in the presence of complete protease inhibitor cocktail (Roche, 1:1000) on ice, followed by 10-20 strokes with a pestle to break the cyst wall. The crude extract containing intact flagella was centrifuged at 10,000 g onto coverslips in a Corex tube with an adaptor in a Beckman JS 13.1 rotor. Fixation was carried out in methanol at −20°C for 5 min, followed by incubation for 1 h at room temperature with the following primary antibodies (all 1:500): mouse anti-α-tubulin, (B-5-1-2 coupled to Alexa-488, Thermo Fisher or DM1A, Sigma), rat anti-tyrosinated tubulin (YL1/2, Merck), mouse anti-acetylated tubulin (T6793; Sigma), mouse anti-Centrin-2 (20H5, Millipore). Secondary antibodies were donkey anti-rabbit conjugated to Alexa 594 (Abcam, 150072) and goat anti-mouse conjugated to Alexa 488 (Thermo Fisher, A11001), both used 1:1000). Indirect immunofluorescence was imaged on LSM700 Zeiss and Leica TCS SP8 STED 3X microscope with a 100 × 1.4 NA oil-immersion objective.

### Transmission electron microscopy and tomography

For chemical fixation (Fig. 2G-H, Fig. 3A, Fig. 4C, Fig. 4E), cells were released from gametocysts by applying pressure on the coverslip mounted on a poly-D-Lysine coated slide and fixed overnight in 2% paraformaldehyde 1% glutaraldehyde in phosphate buffer 0.1 M, pH 7.4 washed in cacodylate buffer (0.1 M, pH 7.4) at 4°C, and post-fixed with 0.5% tannic acid in cacodylate buffer (0.1 M, pH 7.4) for 40 min at room temperature. After two washes in distilled water, cells were post-fixed in 1% osmium tetroxide in cacodylate buffer. Thereafter, samples were washed in distilled water and dehydrated in a graded ethanol series (1x 50%, 1x 70%, 2x 96%, 2x 100%), and finally embedded in epon resin before polymerization overnight at 65°C.

For cryo-fixation (Fig. 3B-C), intact gametocysts bathed in 3% BSA were subjected to high-pressure freezing (HPM100, Leica Microsystems), before freeze-substitution using a temperature-controlled chamber (AFS, Leica Microsystems). The samples were first transferred into plastic tubes containing acetone with 0.2% glutaraldehyde at -90°C for 24 h, and then washed in pure acetone before transfer to 0.1 % tannic acid in acetone for a further 24 h. Next, the temperature was raised to -30°C over the course of 3 days in 2 % osmium, and the samples then transferred to 0.5 % uranyl acetate for 24 h while the temperature rose again from -30°C to -10 °C. The samples were then rinsed in pure acetone three times, the temperature raised to 0°C, and a 50/50 acetone/epon resin mix added. The samples were left still for 2 h before being placed on a rotator at room temperature and increasing concentrations of resin added up to 100%. The samples were then left overnight, before hardening at 65°C for 48 hours.

50 nm sections were cut using a diamond knife on an ultramicrotome (Leica UC7) and collected on single slot copper grids with a formvar support film. Sections were then further stained with lead citrate and uranyl acetate before imaging inside a transmission electron microscope operating at 80 kV (Tecnai Spirit, FEI Company), using a CCD camera (Eagle, FEI Company).

Tilt-series from cryo-fixed sections were acquired on a Tecnai F20 operated at 200 kV (Thermo Fischer Scientific using Thermo Scientific Tomography software in continuous tilt scheme from -60° to +60° in 2° steps at -2.5 µm defocus. Data were recorded with a Falcon III DD camera (Thermo Fisher Scientific) in linear mode at 29’000 × magnification, corresponding to a pixel size of 3.49 Å. Tilt series alignment and tomogram reconstruction was done using EMAN 2.9 [^38^]. 4x binned tomograms with a corresponding pixel size of 13.96 Å were corrected using IsoNet v.0.9 [^39^]. Microtubule and central hub densities were traced in the corrected tomograms using Imod 4.9 [^40^]. Video visualization was done using Chimera 1.14.

### Ultrastructure expansion microscopy

Flagella were isolated and spun onto 10 mm coverslips as above. After methanol fixation, coverslips were incubated overnight at room temperature in an acrylamide/formaldehyde solution (1% AA and 0.7% FA in PBS) under mild agitation. Next, coverslips were incubated in 50 µl monomer solution (19% (wt/wt) SA, 10% (wt/wt) AA, 0.05% (wt/wt) BIS in PBS) supplemented with 0.5% Tetramethylethylenediamine (TEMED) and 0.5% Amonium Persulfate (APS) on a piece of Parafilm for 1 h at 37°C in a moist dark chamber for gelation. All subsequent steps were carried out with mild agitation at room temperature unless otherwise stated. For denaturation, gels were incubated for 15 min in denaturation buffer (200 mM SDS, 200 mM NaCl and 50 mM Tris in distilled water, pH=9) in 5 cm Petri dishes followed by incubation for 1 h on a 95°C hot plate in fresh denaturation buffer. After denaturation, gels were washed extensively with distilled water in 10 cm Petri dishes. Water was exchanged 5 times every 20 min, followed by a wash in distilled water overnight at 4°C. After expansion, the gel size was measured with a ruler to determine the fold expansion, and the gel cut in pieces fitting into a 5 cm Petri dish. Prior to staining, gels were blocked for 1 h in blocking buffer (10mM HEPES (pH=7.4), 3% BSA, 0.1% Tween 20, sodium azide (0.05%)) followed by incubation with rabbit anti-α-tubulin antibodies (Abcam 18251) diluted 1:250 in blocking buffer. Thereafter, gels were again washed three times in blocking buffer for 10 min each, before incubation with secondary antibodies diluted in blocking buffer at 37°C in the dark for 3 h. Finally, gels were washed three times in blocking buffer for 10 min each before transfer into a 10 cm Petri dish for re-expansion by 6 washes, each 20 min in distilled water. For imaging, gels were cut and mounted on a 60×24 mm coverslip coated with poly-D-lysine diluted in water (2 mg/ml) and supported on both longitudinal sides with capillaries attached with superglue. To prevent drying, the edges of the gel were covered with VaLaP and the gel covered with Halocarbon oil 700 for imaging. STED images were acquired using the 775 pulsed laser for depletion on a Leica TCS SP8 STED 3X microscope with a 100 × 1.4 NA oil-immersion objective.

### Image analysis

For quantification of axoneme and centriole width in α-tubulin UExM STED experiments, three 1 µm line scans with a width of 5 pixel perpendicular to the long axis of flagellum were measured per image. Peak calling in IgorPro8 software (Wavemetrics, USA) was used to first interpolate and then resample to obtain the same number of data points for all images regardless of the expansion factor. Further line profile peak analysis was done using its build-in PeakFind function. The parameters were set globally for all traces as follows: pBegin 30; pEnd 100; maxPeaks 6; minPeakPercent 5; noiselevel 1.4; smoothingfactor 5.2. The most frequent categories remained unchanged if the threshold was altered. Next, mean intensity line profiles for both the axoneme and the centriole were calculated, and full-width half maxima (FWHM) determined using a custom Python script, and the average of the FWHMs computed. Line profiles were aligned on the center of the FWHM to plot the mean intensity profile from all 18 profiles combined, together with the standard deviation.

In EM and UExM transverse views, centrioles with slight perspective distortion due to tilted orientations were circularized using the ImageJ plugin ‘Transform-Interactive Affine’ before symmetrization. The circularized input image and the symmetrized images from 4- to 11-fold symmetry were overlaid and both intensity profiles measured in a circular band along the microtubules, and the correlation of the signal analyzed using R2 values for each symmetrization.

The ‘angle tool’ in FIJI was used for quantification of angles in wedges between the center of the centriole and two clearly visible neighboring peripheral A-microtubules in centrioles and axonemes imaged in transverse view by EM.

## References

1. Gönczy, P. & Hatzopoulos, G. N. Centriole assembly at a glance. J. Cell Sci. 132, jcs228833 (2019).

2. Breslow, D. K. & Holland, A. J. Mechanism and Regulation of Centriole and Cilium Biogenesis. Annu. Rev. Biochem. 88, 691–724 (2019).

3. Azimzadeh, J.Evolution of the centrosome, from the periphery to the center. Curr. Opin. Struct. Biol. 66, 96–103 (2021).

4. Jana, S. C. et al. Differential regulation of transition zone and centriole proteins contributes to ciliary base diversity. Nat. Cell Biol. 20, 928–941 (2018).

5. Loreng, T. D. & Smith, E. F. The Central Apparatus of Cilia and Eukaryotic Flagella. Cold Spring Harb. Perspect. Biol. 9, a028118 (2017).

6. Karpov, S. A. et al. The Chytrid-like Parasites of Algae Amoeboradix gromovi gen. et sp. nov. and Sanchytrium tribonematis Belong to a New Fungal Lineage. Protist 169, 122–140 (2018).

7. Karpov, S. A., Vishnyakov, A. E., Moreira, D. & López-García, P. The Ultrastructure of Sanchytrium tribonematis (Sanchytriaceae, Fungi incertae sedis) Confirms its Close Relationship to Amoeboradix. J. Eukaryot. Microbiol. 66, 892–898 (2019).

8. Sinden, R., Talman, A., Marques, S., Wass, M. & Sternberg, M. The flagellum in malarial parasites. Curr. Opin. Microbiol. 13, 491–500 (2010).

9. Dubremetz, J. F. & Yvore, P. [Oocystic wall formation in the coccidia Eimeria necatrix Johnson 1930 (Sporozoa, Coccidiomorpha). Study with electronic microscope]. C. R. Seances Soc. Biol. Fil. 165, 862–866 (1971).

10. Pelletier, L., O’Toole, E., Schwager, A., Hyman, A. A. & Müller-Reichert, T. Centriole assembly in Caenorhabditis elegans. Nature 444, 619–623 (2006).

11. Roggen, D. R., Raski, D. J. & Jones, N. O. Cilia in Nematode Sensory Organs. Science 152, 515–516 (1966).

12. Ross, M. M. Modified cilia in sensory organs of juvenile stages of a parasitic nematode. Science 156, 1494–1495 (1967).

13. Dallai, R., Xué, L. & Yin, W. Flagellate spermatozoa of Protura (Insecta, Apterygota) are motile. Int. J. Insect Morphol. Embryol. 21, 137–148 (1992).

14. van Deurs, B. Axonemal 12 + 0 pattern in the flagellum of the motile spermatozoon of Nymphon leptocheles. J. Ultrastruct. Res. 42, 594–598 (1973).

15. van Deurs, B. Pycnogonid sperm. An example of inter- and intraspecific axonemal variation. Cell Tissue Res. 149, 105–111 (1974).

16. King, P. E. & El-Hawawi, A. S. N. Spermiogenesis in the Pycnogonid Pycnogonum littorale (Ström). Acta Zool. 59, 97–103 (1978).

17. Baccetti, B., Dallai, R. & Fratello, B. The spermatozoon of arthropoda. XXII. The 12+0’, 14+0’ or aflagellate sperm of protura. J. Cell Sci. 13, 321–335 (1973).

18. Phillips, D. M. FINE STRUCTURE OF SCIARA COPROPHILA SPERM. J. Cell Biol. 30, 499–517 (1966).

19. Phillips, D. M. OBSERVATIONS ON SPERMIOGENESIS IN THE FUNGUS GNAT SCIARA COPROPHILA. J. Cell Biol. 30, 477–497 (1966).

20. Reger, J.F. and Florendo, N.T. Observations on microgamonts and microgametes of the coccidian, Eimeria sp. parasitic in the ostracod, Cypridopsis sp. J. Submicrosc. Cytol 2, 69–78 (1970).

21. Prensier, G., Vivier, E., Goldstein, S. & Schrével, J. Motile Flagellum with a ‘3 + 0’ Ultrastructure. Science 207, 1493–1494 (1980).

22. Schrevel, J. & Besse, C. [A functional flagella with a 6 + 0 pattern]. J. Cell Biol. 66, 492–507 (1975).

23. Schrével, J. Recherches sur le cycle des Lecudinidae grégarines parasites d’annélides polychétes. Protistologica 5, 561–588. (1969).

24. Francia, M. E., Dubremetz, J.-F. & Morrissette, N. S. Basal body structure and composition in the apicomplexans Toxoplasma and Plasmodium. Cilia 5, 3 (2015).

25. Kuriyama, R., Besse, C., Gèze, M., Omoto, C. K. & Schrével, J. Dynamic organization of microtubules and microtubule-organizing centers during the sexual phase of a parasitic protozoan,Lecudina tuzetae (Gregarine, Apicomplexa). Cell Motil. Cytoskeleton 62, 195–209 (2005).

26. Friedman, M. H. A reevaluation of the Markham rotation technique using model systems. J. Ultrastruct. Res. 32, 226–236 (1970).

27. Guichard, P. et al. Native architecture of the centriole proximal region reveals features underlying its 9-fold radial symmetry. Curr. Biol. CB 23, 1620–1628 (2013).

28. Chen, F., Tillberg, P. W. & Boyden, E. S. Optical imaging. Expansion microscopy. Science 347, 543–548 (2015).

29. Gambarotto, D. et al. Imaging cellular ultrastructures using expansion microscopy (U-ExM). Nat. Methods 16, 71–74 (2019).

30. Nazarov, S. et al. Novel features of centriole polarity and cartwheel stacking revealed by cryo-tomography. EMBO J. 39, e106249 (2020).

31. Gönczy, P. Towards a molecular architecture of centriole assembly. Nat. Rev. Mol. Cell Biol. 13, 425–435 (2012).

32. Hirono, M. Cartwheel assembly. Philos. Trans. R. Soc. Lond. B. Biol. Sci. 369, 20130458 (2014).

33. Hilbert, M. et al. SAS-6 engineering reveals interdependence between cartwheel and microtubules in determining centriole architecture. Nat. Cell Biol. 18, 393–403 (2016).

34. Banterle, N. et al. Kinetic and structural roles for the surface in guiding SAS-6 self-assembly to direct centriole architecture. Nat. Commun. 12, 6180 (2021).

35. Banterle, N. & Gönczy, P. Centriole Biogenesis: From Identifying the Characters to Understanding the Plot. Annu. Rev. Cell Dev. Biol. 33, 23– 49 (2017).

36. Akella, J. S. et al. Cell type-specific structural plasticity of the ciliary transition zone in C. elegans. Biol. Cell 111, 95–107 (2019).

37. Goldstein, S. F. Motility of 9 + 0 mutants of Chlamydomonas rheinhardtii. Prog. Clin. Biol. Res. 80, 165–168 (1982).

38. Tang, G. et al. EMAN2: an extensible image processing suite for electron microscopy. J. Struct. Biol. 157, 38–46 (2007).

39. Liu, Y.-T. et al. Isotropic Reconstruction of Electron Tomograms with Deep Learning. http://biorxiv.org/lookup/doi/10.1101/2021.07.17.452128 (2021) doi:10.1101/2021.07.17.452128.

40. Kremer, J. R., Mastronarde, D. N. & McIntosh, J. R. Computer visualization of three-dimensional image data using IMOD. J. Struct. Biol. 116, 71–76 (1996).

